# Putting a new spin on insect jumping performance using 3D modeling and computer simulations of spotted lanternfly nymphs

**DOI:** 10.1101/2023.06.20.545794

**Authors:** Chengpei Li, Aaron J. Xu, Eric Beery, S. Tonia Hsieh, Suzanne Amador Kane

## Abstract

How animals jump and land on a variety of surfaces is an ecologically important problem relevant to bioinspired robotics. We investigated this topic in the context of the jumping biomechanics of the planthopper *Lycorma delicatula* (the spotted lanternfly, SLF), an invasive insect in the US that jumps frequently for dispersal, locomotion, and predator evasion. High-speed video was used to analyze jumping by SLF nymphs from take-off to impact on compliant surfaces. These insects used rapid hindleg extensions to achieve high take-off speeds (2.7-3.4 m/s) and accelerations (800-1000 ms^-2^), with midair trajectories consistent with zero-drag ballistic motion without steering. Despite rotating rapidly (5-45 Hz) in the air about time-varying axes of rotation, they landed successfully in 58.9% of trials; they also attained the most successful impact orientation significantly more often than predicted by chance, consistent with their using attitude control. Notably, these insects were able to land successfully when impacting surfaces at all angles, pointing to the emerging importance of collisional recovery behaviors. To further understand their rotational dynamics, we created realistic 3D rendered models of SLFs and used them to compute their mechanical properties during jumping. Computer simulations based on these models and drag torques estimated from fits to tracked data successfully predicted several features of their measured rotational kinematics. This analysis showed that SLF nymphs are able to use posture changes and drag torques to control their angular velocity, and hence their orientation, thereby facilitating predominately successful landings when jumping.

**Summary:** High-speed video revealed that juvenile spotted lanternflies are adept at landing after tumbling rapidly midair during jumping. We present computer simulations and realistic 3D models to help explain these abilities.

## Introduction

Jumping is an energetically costly mode of locomotion, requiring large force generation and challenging levels of sensorimotor control (Biewener and Patek, 2018). Animals jump to escape predators, capture prey and gain access to resources, traverse complex environments, and navigate obstacles. Because jumping can be critical for survival, it is employed among a wide diversity of animal taxa across a broad range of sizes (Hawkes et al., 2022; Mo et al., 2020b). Biomechanical studies that determine how animals accomplish such demanding feats are therefore both ecologically relevant and a significant source of inspiration for the design of jumping robots (Mo et al., 2020b; Ribak, 2020). Motions during jumping can be broken into three phases: take-off, midair, and impact and landing. Most prior studies have focused on the initial take-off period most critical for achieving jumps with optimal values of height and horizontal range (e.g., see (Burrows et al., 2021; Mo et al., 2020b) and references therein). However, jumping also presents the organism with challenges in controlling both the direction of its jump and its body orientation throughout the trajectory. Therefore, some recent studies have considered how the midair phase is not simply a passive period of parabolic flight, but rather can include active and passive stabilization measures, as well as directed navigation critical to a controlled landing (Burrows et al., 2015; Ortega-Jimenez et al., 2022; Zong et al., 2022)).

Among insects, planthoppers (infraorder Fulgoromorpha) in particular have exceptional jumping abilities that have been studied extensively in the preparatory and take-off phases (Burrows, 2009; Burrows, 2014a; Burrows et al., 2019). Although these insects make extensive use of jumping, it is less well-documented how they manage the midair and landing phases. The spotted lanternfly (*Lycorma delicatula*) (SLF) is a species of planthopper that has emerged recently as a highly-invasive pest in the US and South Korea, posing a significant agricultural threat. SLFs jump as a dominant form of locomotion to disperse, find food and habitat, and evade predators (Nixon et al., 2021). Like many other insects, they also need to be able to land on complex landscapes formed by densely-clustered leaves oriented at random, unpredictable angles (Graham and Socha, 2020; Kane et al., 2021; Zeng et al., 2020). In this study, we present the first exploration of SLF jumping biomechanics using an interdisciplinary approach that utilizes both empirical data on jumping kinematics using high-speed 3D video, detailed 3D rendered models, and computer simulations of the resulting trajectories and rotational dynamics. In addition to characterizing their jumping performance at take-off, we also considered how these insects negotiate the post-take-off midair and landing phases. We hypothesized that SLFs would extend their legs during the midair phase of a jump to decrease the body’s angular speed and increase the probability of a successful landing.

In particular, we studied whether SLF nymphs utilize several midair behaviors observed for other insects. First, we considered directed aerial descent, in which jumping and falling insects and spiders change their trajectory direction in midair (Yanoviak et al., 2010; Yanoviak et al., 2011; Yanoviak et al., 2015; Zeng et al., 2015) to enhance their ability to land on an intended target (Socha et al., 2015). To test the hypothesis that SLF nymphs might use directed aerial descent, we measured whether their tracks while in the air deviated from a planar ballistic trajectory, and whether the timing of changes in their posture correlated with changes in body orientation, heading, speed, and angular velocity (Yanoviak et al., 2010; Yanoviak et al., 2015).

Second, we studied whether SLF nymphs can reorient their bodies while in the air (attitude control), to facilitate landing. Such aerial reorientation can be a passive consequence of aerodynamic drag and body posture (e.g., dragonflies (Fabian et al., 2021), pea aphids (Ribak et al., 2013), spiders (Yanoviak et al., 2015), SLF nymphs (Kane et al., 2021), and springtails (Ortega-Jimenez et al., 2022)), or can involve active body motions (e.g., stick insect nymphs (Zeng et al., 2017), mantises (Burrows et al., 2015), lizards (Higham et al., 2017; Jusufi et al., 2008; Siddall et al., 2021), frogs (Wang et al., 2022), and squirrels (Fukushima et al., 2021)). We therefore studied whether and how SLF nymphs attain different body postures at different phases during jumping, particularly before landing attempts.

Third, we investigated whether SLF nymphs rotate rapidly after take-off, as has been observed for several other insect species (Brackenbury, 1996; Burrows, 2012; Burrows et al., 2007; Burrows et al., 2015; Goode and Sutton, 2023; Zong et al., 2022). Such behaviors are ecologically salient because midair rotational motion can complicate self-righting, targeting and landing, as well as potentially generate lift and thereby extend the aerial trajectory via the Magnus effect (Wauthy et al., 1998), and enhance predator evasion (Burrows and Dorosenko, 2014). Several recent studies have explored how spinning insects can reduce their angular velocity and reorient in midair to facilitate landing upright by bending their flexible bodies (Ortega-Jimenez et al., 2022)) or deploying their wings (Zong et al., 2022). To determine whether SLF nymphs indeed rotate during jumping, and whether they use changes in body posture or drag torques to control their angular velocity and orientation, we used video tracking data to measure their rotational motion during jumping, and to test for correlations between angular velocity and changes in body posture. To interpret this video data, we expanded on earlier studies that modeled elongated insects as assemblies of jointed rods (Burrows et al., 2015; Zeng et al., 2017) by creating detailed 3D rendered models of the insect’s body in postures observed during various jumping phases, and used these to parameterize computer simulations of their rotational dynamics.

## Methods

### Insect methods

Third and fourth instar SLF nymphs (Fig. 1A) were collected between mid-morning and late afternoon (Kim et al., 2011) from *Ailanthus altissima* (tree-of-heaven) and *Vitis vinifera* (wild grape vines) in southeastern Pennsylvania (40°00’ 30.2” N, 75°18’ 22.0” W) in late June through early July 2021. Specimens were maintained on the leaves of potted ≥ 1 m tall *A. altissima*, their preferred native host plant, as recommended for raising SLFs in the laboratory (Nixon et al., 2022), and used for experiments within 2 hours of collection. For this purpose, *A. altissima* trees were collected from the field with intact root systems, maintained in mesh cages under full-spectrum grow lights, and replaced before they began to show stress from SLF feeding. After experiments were completed, the body mass of each specimen was measured and used to determine life stage for the third instars (Bien et al., 2023). Laboratory environmental conditions were maintained at 23 [22, 24] deg C and 64 [44, 73] % relative humidity.

**Figure 1.**
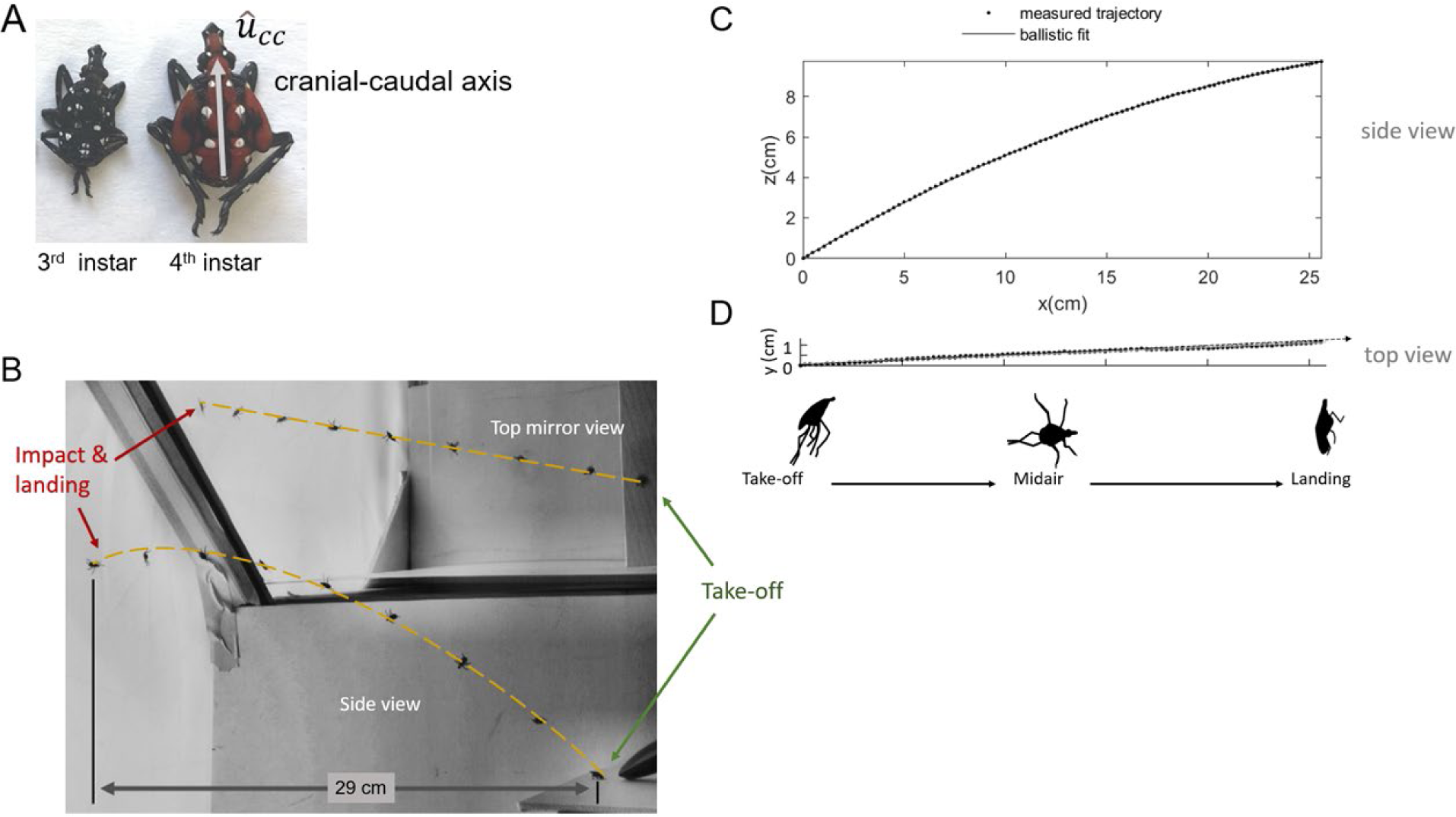
A) Spotted lanternfly nymph 3^rd^ and 4^th^ instars. Gray arrow: cranial-caudal axis, *û_cc_*, tracked to measure body orientation vs time. (Adapted from (Bien et al., 2023).) B) Jumping experiment arena photograph showing the launch pad (bottom left), superimposed images of a spotted lanternfly nymph at different times and the jumping trajectories (dashed orange line), and the white fabric target (far right) on which they impacted and landed, as seen in the main camera side view and orthogonal top mirror view. Spotted lanternfly nymph jumping raw trajectory coordinate data (symbols) and 3D ballistic fits (lines) as seen from the C) side and D) top views. Here, *z* is the vertical, *x* and *y* are in the horizontal plane.

### Data analysis and statistics

All fitting and statistical analyses were performed in MATLAB vR2022a (Mathworks, Natick MA USA) and R (R Core Team, 2017); all mentions of MATLAB function calls are italicized. Hypothesis testing was performed using a significance level of 0.05. Data are reported as grand means [95% CI] computed from means of individual specimen means, unless stated otherwise.

### Jumping experiments

Two preliminary field studies were performed to aid in the design of laboratory experiments. To determine whether jumping behaviors in the laboratory resembled those in the field, SLF 4^th^ instar nymphs were filmed jumping and landing on leaves, either spontaneously or in response to being prodded by a finger, using a GoPro Hero 9 Black Edition (1920 ×1080 pixels, 240 frame/s); only six videos of six different individuals were obtained due to the challenges inherent in filming small insects in dense foliage. In addition, to check for fatigue due to repeated jumping, we observed 4^th^ instar SLF nymphs (N = 6) that were placed a stone walkway in the shade and then stimulated to jump toward the sunlit bed of foliage upon which they were collected. The starting and end position of each jump was marked and their distance measured to determine jumping ranges. Linear regression performed on the data found weak correlation between jumping range (85 ± 16 mean ± s.e.m.) and jump number for all specimens (R-squared ≤ 0.22), with only one of the six specimens having a significant dependence between range and jump number (p = 0.042). (Fig. S1, Table S1) This supported studying multiple jumping trials by individual specimens in the laboratory experiments.

In the laboratory, we recorded high-speed 3D video for SLF nymphs jumping between a raised platform to a compliant vertical target, a geometry chosen to resemble jumping between different locations in plants while providing an unimpeded view of the insect. The horizontal 16.5 deep × 20.5 wide x 23 cm high take-off platform was made of 80-grit sandpaper spray-painted white and mounted on hardboard; this rough substrate was used because the specimens avoided jumping from smoother surfaces (e.g., posterboard or glass), instead electing to walk away or climb to the underside of the platform. The jumping experiment arena (Fig. 1B) had white acrylic and posterboard walls on two sides, a clear plexiglass wall for observing and positioning specimens, and white insect netting to reduce air currents and prevent escapes. Previous research indicated this species exhibits positive phototaxis (Domingue and Baker, 2019; Jang et al., 2013), so specimens were released in front of a brightly-backlit white mesh insect netting placed over the white fabric cover of the LED light source used to illuminate the scene for videography; the netting also served as a compliant surface that the nymphs were able to grasp and cling to. The horizontal distance from the take-off platform to the target was approximately 29 cm, a value chosen as approximately half the mean horizontal jumping ranges found for 3^rd^ and 4^th^ instar SLF nymphs (Nixon et al., 2021).

Videos were filmed using a single Chronos 1.4 video camera (1280 × 1024 pixel; 1000 fps, 200 microsec exposure time) that captured the trajectory in two orthogonal views: a side view in the approximate plane of the trajectory, and an overhead view reflected in a mirror oriented at 45 deg to the horizontal (image resolution: 3.6 and 3.0 pixel/mm respectively). The arena was illuminated by a Godox white LED lamp at intensities (22,100 ± 100 lux) similar to values measured in the open areas surrounding trees where SLFs were collected.

To induce SLF nymphs to jump, specimens were placed gently onto the take-off platform using scoop-shaped forceps and allowed to move freely. If a nymph oriented toward the target, it was encouraged to jump by approaching it rapidly with a finger or pen to simulate predator attack. The jumping strategies explored here therefore likely reflect escape behaviors. When possible, specimens were recaptured and induced to jump for repeated trials in the same session.

Experiments were performed with the goal of obtaining at least 15 videos for the two nymphal life stages studied that showed the entire jump trajectory in both camera views from take-off to either successful or attempted and failed landing (“analyzable” videos). All videos recorded were viewed repeatedly to find those that matched these criteria and then analyzed for jumping performance. Videos that showed at least the body orientation at impact and the outcome (successful or failed landing attempts) were separately scored to study behavior during attempted landing.

### Video analysis and trajectory fitting

All analyzable videos were scored manually to determine the postures assumed during jumping, the time required to assume each posture, time of first impact, landing orientation and strategy, and whether the specimen landed successfully. The phases during jumping were defined as take-off (when leg motions were first evident to when the feet first lost contact with the surface), midair (take-off to first impact), impact (when the body first contacted a surface) and landing. A successful landing was defined as an impact in which the SLF nymph was able to cling to the surface, and a failed landing as one in which the SLF nymph either bounced off or fell off of the target surface. The orientation at impact was estimated from the closest match between the landing surface and the body ventral, dorsal, right, left, cranial or caudal surfaces. To test for dependence of landing success on orientation at impact, we compared our measured proportions to a simple model of equal probabilities (1/6) of landing in each of the six orientations using MultinomCI (R Core Team, 2017) to compute the 95% CI for the measured proportions for each outcome.

The SLF nymph body center of mass (COM) was tracked throughout each trial on both camera views using a combination of custom MATLAB code and semi-automatic tracking using the program DLTdv (Hedrick, 2008). A combination of DLTdv and 3D reconstruction parameters determined by wand calibration (Theriault et al., 2014) (wand score of 0.6, wand end point s.d. = 0.4 mm (0.4% span), and pixel projection errors 0.11-0.15 pixels ≤ 0.54 mm) was used to convert the 2D tracked coordinates on the two camera views into 3D coordinates with the origin at the point of take-off, *z* oriented vertically upward (as determined by a freely-falling ball), and *x* and *y* in the horizontal plane. (Fig. 1C) The tracked coordinates vs time were used to compute the velocity direction (heading) using quadratic polynomial regression over 25 ms time windows; this method resulted in a root-mean-square-error between the fit and measured track data equal to the 3D reconstruction uncertainty.

The raw 3D trajectories from take-off to impact were fitted to a zero-drag ballistic model using nonlinear least squares regression, an approach used to characterize the kinematics of jumping performance of spiders and crickets (Nabawy et al., 2018; Palmer et al., 2018), using the equation for a parabola:

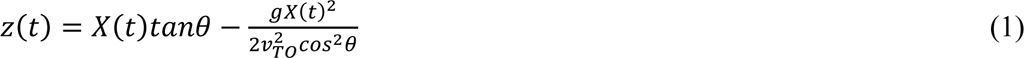

where *X*(*t*) is distance from the position at take-off projected onto the horizontal x-y plane (Fig. 1C), *v_TO_* is take-off speed, *g* = 9.81 m/s^2^ is the acceleration of gravity, and *θ* is take-off angle with respect to the horizontal (Taylor, 2005).

We used the results of fitting to look for three effects that can cause the measured trajectories to deviate from the simple zero-drag ballistic model. First, drag forces can cause the trajectory to deviate systematically from a parabola at longer times. Second, body rotations and changes in posture can cause the trajectory of the whole insect COM to deviate from that for the body-only COM, which was tracked on video; this can result in nonrandom variations in the fit residuals and heading angles. Third, while a zero-drag ballistic trajectory lies in a vertical plane, steering can cause variations in the azimuthal heading angle (direction of the velocity’s horizontal component), and hence a nonplanar trajectory. To test for steering, the 3D coordinates of each trajectory were also fit to a plane using *pcfitplane*. To test for these three effects, we assessed the goodness-of-fit of the zero-drag, no-steering model by comparing the residuals for the ballistic trajectory fit, *e_ballistic_*, (Eqn 1) and the planar fit, *e_planar_* to the 3D reconstruction uncertainty. Plots of these fit residuals vs time also were examined for nonrandom variation by comparing the rms fit residuals between three time intervals: 1) within 25 ms after take-off; 2) within 25 ms after midair posture changes, and 3) within 5 ms of impact.

Jumping performance was characterized by computing the acceleration during take-off using *a*_TO_ = *v*_TO_/*T_accel_*. The maximum jumping range, *R*, and time-of-flight, *T_TOF_*, corresponding to fitted values of *v_TO_* and the optimal take-off angle *θ* = 45 deg, were estimated using *R* = *v_TO_* _2_ sin^2^ *θ* /*g* and *T_TOF_* = 2 *v_TO_* sin *θ* /*g* (Taylor, 2005). We also characterized the aerodynamic regime during jumping by computing the Reynolds number, *Re = ρ v_TO_ L / μ,* where air density, *ρ* = 1.2 kg m^-3^, body length, *L_b_* = 9.3 and 11.7 mm for 3^rd^ and 4^th^ instar SLF nymphs, respectively, and the dynamic viscosity of air, *μ* = 18.3 × 10^-6^ Pa-s (Alexander, 2016).

### Measuring and simulating rotational kinematics and dynamics

SLF nymph body orientation and rotational kinematics during jumping was tracked manually on video in 3D using DLTdv; results are reported relative to a coordinate frame, *S_s_* (the “spatial frame”) with its axis directions fixed relative to the laboratory and its origin at the nymph’s COM. SLF nymph angular orientation and velocity were described using the Tait-Bryan (roll-pitch-yaw) angle convention in ZYX order, using the orientation of the body principal axes of rotations defined below relative to the xyz axes in the laboratory defined above. The initial 3D angular velocity, ***ω***(0), was computing using the MATLAB function *angvel* with the 3D orientation of the nymph immediately after take-off and 5 ms afterward. Nymphs were not resolved well enough on video for the entire jumping trajectory to allow tracking of their full 3D orientation on every frame. Instead, we tracked the orientation of their cranial-caudal axis, **û_cc_**, for the entire trajectory. (Fig. 1A) This was done by tracking one point on the head and one on the caudal end on each frame from initial take-off until impact on the landing target, and then using custom MATLAB code to compute the angular speed using: *ω_cc_*(*t*) = ***u_cc_***/Δ*t*|. The values of *ω_cc_* vs time were used to compute several measures of rotational kinematics, including angular speed at take-off, *ω_TO_*, and the ratio of angular speed at leg extension to that at take-off, *ω_ext_ / ω_TO,_*. Because the tracked data for some trials exhibited oscillations in *ω_cc_(t)* vs time, the oscillation period, *Tosc*, was measured manually from the time intervals between successive peaks. For consistency with earlier studies, we report ω in units of Hz, where 1 Hz = 2π rad/s.

Computer simulations were used to reconstruct full 3D rotations and interpret how the midair rotational motions during jumping related to the observed SLF nymph body postures. This allowed us to test hypotheses about whether these insects use attitude control. The 3D rotational dynamics during jumping was modeled using the following results from advanced classical mechanics; see (Goldstein, 1980) for an in-depth treatment. The equation of motion for the simplest case of a point mass, *m*, rotating about a fixed axis is:

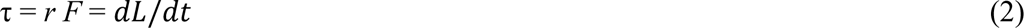

Here τ is the net external torque (moment of force), *r* is the magnitude of the moment arm (the vector from the point mass to the axis of rotation), *F* is the component of force perpendicular to the moment arm, *t* is time, and *L* is angular momentum, *L* = *I_rot_ ω*, where *I_rot_* = mr^2^ is the moment of inertia, a measure of resistance to changes in angular velocity, and *ω* is angular velocity in rad/s (Taylor, 2005). (Fig. 2A) For an extended rigid object rotating about a given axis, *I_rot_* depends on the object’s mass distribution relative to the axis of rotation (Fig. 2B) as:

**Figure 2.**
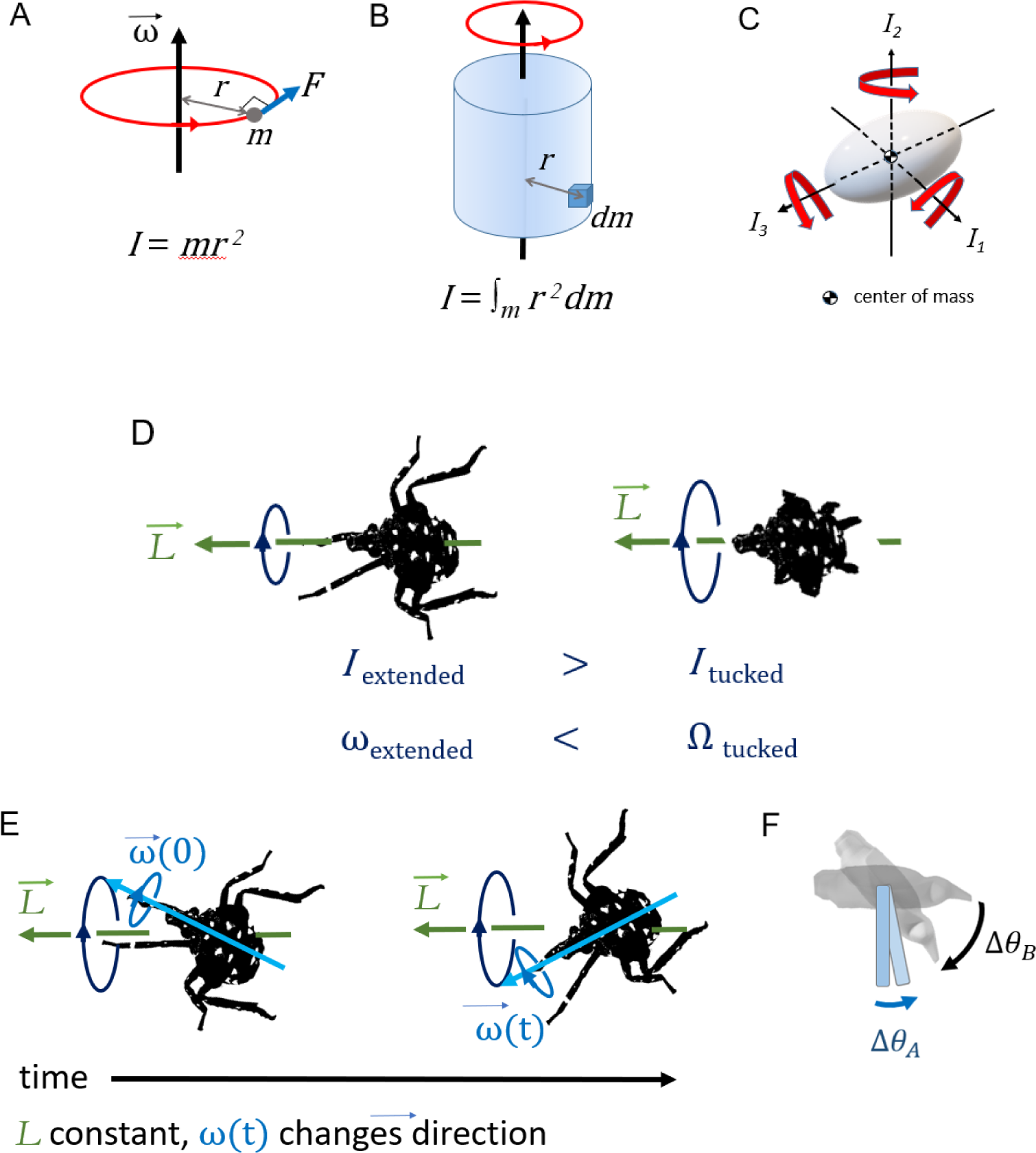
A) Illustration of the geometry used in defining the quantities used to describe rotational dynamics about a single axis for a point mass. B) Geometry for computing moment of inertia, *I*, for an extended rigid body. C) Schematic illustration of the principal axes of rotation for an extended body, labeled by their corresponding values of moment of inertia. D) For zero external torque, angular momentum, *L*, is constant, so changes in body posture that alter moment of inertia, *I_rot_*, also change angular velocity, ω. E) Illustration of the geometry for torque-free precession, in which the rotational velocity vector, ω^→^ and its associated axis of rotation (blue arrows) describes a cone about the constant angular momentum vector, *L*^→^ (green arrow) even if the external torque is zero. F) Illustration of a simple model with two-jointed rods to illustrate the geometry for Eqn 7 and the surrounding discussion.

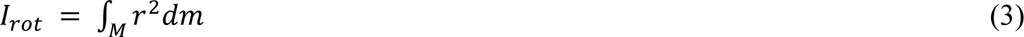

In 3D, torque and angular velocity are both vectors and the moment of inertia is a 3×3 matrix, the inertia tensor, ***I_rot_*** (Goldstein, 1980). There exists a coordinate frame, *S_o_* (the “object frame”) with its origin at the COM in which the object’s orientation is constant and the inertia tensor is a diagonal matrix with elements *I_1_*, *I_2_*, and *I_3_*. In this frame, the coordinate axes, *û_i_*, are called the object’s principal axes, and *I_i_* is the moment of inertia for rotations about the *i*th principal axis. (Fig. 2C)

The full 3D rotational dynamics of a rigid body in frame *S_o_* with its origin fixed at the object’s COM are given by the Euler equations (Goldstein, 1980):

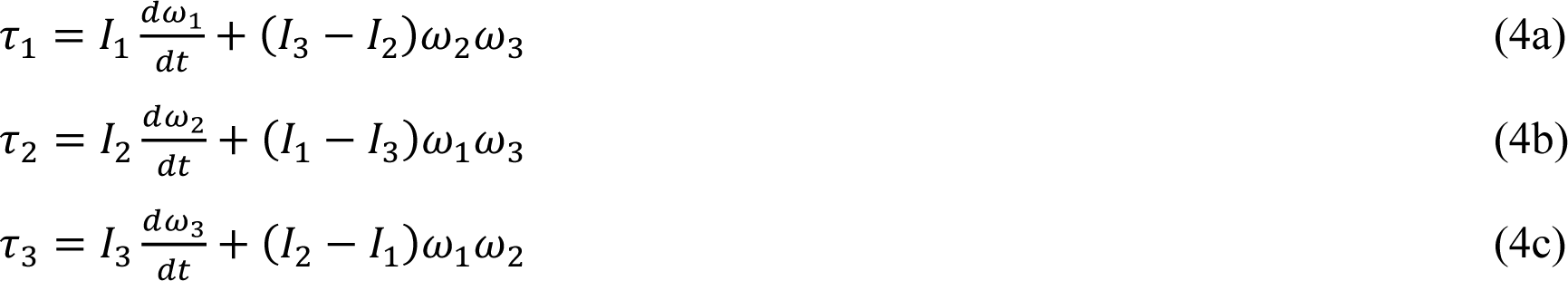

These equations can be integrated to solve for the object’s angular velocity vector, ***ω***(*t*) = (ω_1_, ω_2_, ω_3_), given *I_1_*, *I_2_*, and *I_3_* and the initial angular velocity, ***ω***(0) and torque vector in frame *S_o_*. We solved Eqn 4 using MATLAB’s ordinary differential equation solver, *ode45* using a simulation timestep 2.6×10^-6^ s (≤ 0.04 deg/timestep), using ***ω***(0) from tracking and torque and the inertial tensor estimated as described below (Marghitu and Dupac, 2012). The angular velocity for each timestep then was converted into a sequential series of 3D rotations to give the object’s rotational motion in the spatial frame, *S_s_*, using custom MATLAB code.

During jumping, after take-off the only external torque is due to aerodynamic drag, *τ*_*drag*_, which is related to angular velocity along each principal axis by:

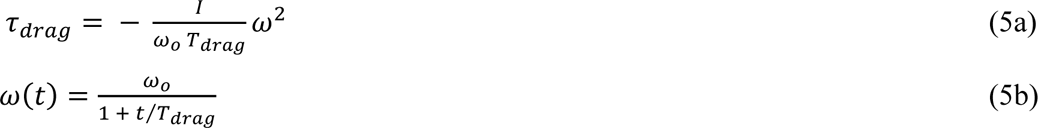

where *T*_*drag*_ is a characteristic time for the angular velocity to decrease to half its initial value, ω*_o_* = |***ω***(0)| (Taylor, 2005). Measured values of *ω_cc_(t)* from tracking were fitted to Eqn 5b using nonlinear least squares fits in MATLAB to find *T*_*drag*_; this was then used with Eqn 5a to estimate the torques used in solving Eqn 4 to model the influence of aerodynamic torques on SLF nymph rotations.

Video analysis showed that SLF nymphs assume characteristic body postures during jumping, and that the axis of rotation often changed during the midair phase even for a fixed posture. To explain this phenomenon, we note that the Euler equations allow the direction of the angular velocity to change with time even if net external torque is zero and angular momentum is constant because in 3D, angular momentum is given by ***L* = *I_rot_ · ω***. If the insect’s body has 2 or more 3 different values of *I_i_*, this means that ***L*** and ***ω*** only have the same direction for rotations purely about one principal axis. If an insect with constant posture does not rotate about a principal axis, it will undergo a form of rotational motion called torque-free precession in which its angular velocity describes a cone about the constant angular momentum vector, (Butikov, 2006). (Fig. 2E) We therefore computed the values of *I_i_* for the observed nymph postures to test if this argument (i.e., unequal values of *I_i_*) could account for their observed rotations.

We also modeled how the observed changes in posture by SLF nymphs affect their moments of inertia, and hence body orientation and angular velocity as a function of time. One possibility is that they might extend or tuck their legs, similar to when a human tucks or extends their arms and legs during a dive, so as to keep the principal axes constant but change one or more values of *I_i_*. (Fig. 2D) If drag torque is negligible, then in this scenario Eqn 4 reduces to *L_i_* = *I*_*i*_*ω_i_* = constant (angular momentum conservation) along each principal axis, and the moments of inertia and angular velocity for the two postures then are related as:

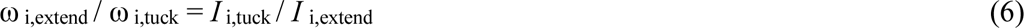

Eqn 6 explains how divers can spin faster by tucking their arms and legs (increasing ω _i_ by decreasing *I*_*i*_) or spin more slowly by extending them (decreasing ω _i_ by increasing *I*_*i*_). More generally, changes in body posture can alter the body’s orientation and rotational axis, not only its angular speed. For example, this occurs when a diver “throws” one or more appendages (i.e., rapidly rotates arms or legs across the body) during a dive. If this change in posture also reorients the principal axes, it will change the direction of ***ω*** even though angular momentum is constant. This can be understood using an articulated model in which the body (B) and appendages can rotate relative to each other, such that total angular momentum, ***L* = *L*_B_+ *L*_A_**. The throw can be interpreted as exchanging angular momentum between the body and appendages (A) such that the change in ***L*, Δ*L***, is zero and hence Δ***L*_B_= - Δ*L*_A_**. For this reason, the resulting motion is called an angular momentum twist. Competition divers use this effect to execute twisting dives in which, e.g., they initially rotate by pitching, perform a throw to execute a complete roll, then execute a second throw so as to resume pitching (Frohlich, 1979). Sequential application of this mechanism also allow animals to achieve a preferred orientation midair: e.g., falling cats can use it to self-right aerially (see (Frohlich, 1980) for a non-mathematical discussion). A simple model of this can be constructed by considering that the object consists of two jointed rigid segments with moment of inertia *I*_*A*_ and *I*_*B*_ oriented by a relative angle *θ* along a principal axis. If segment A rotates by Δ*θ_A_* over time *dt*, then B must counterrotate to conserve angular momentum, resulting in a body rotation of Δ*θ_B_* (Fig. 2F)

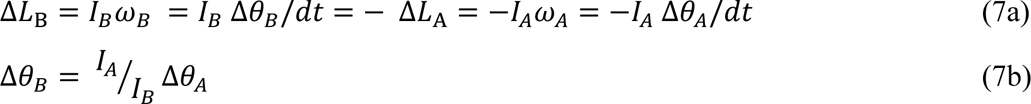

After this maneuver, the insect can maintain this body rotation, either by keeping the legs fixed or tucking them close to the body without changing angular momentum. Previous studies have combined video tracking and modeling to show how angular momentum exchange influences aerial reorientation by insects with rod-like bodies and legs, jumping mantises and falling stick insect instars (Burrows et al., 2015; Zeng et al., 2017). Here we modeled how the observed changes in SLF nymph posture during jumping affect both *I*_*i*_ and the principal axis direction, and therefore the rotational motions predicted by Eqn 4.

### 3D modeling and calculations of mechanical properties

Realistic 3D models of SLF nymphs were created for calculating their mechanical properties for use in computer simulations. First, reference images were recorded using freshly-euthanized SLF nymphs mounted on a size #00 insect pin and photographed from multiple angles using a Sony DSC-RX100 camera in macro mode under diffuse LED lighting. Photographs were obtained at 5 different camera elevation angles for every 10 deg of horizontal rotation using a turntable to rotate the specimen. The resulting photos were then analyzed by photogrammetry using Meshroom, an open-source 3D reconstruction program (https://alicevision.org/#meshroom, 1/16/2023) to create a 3D mesh of the photographed surface. (Fig. 3A) Following methods outlined in (Semple et al., 2019), the 3D mesh and the measured body and leg dimensions were used with the open-source 3D graphics package Blender (www.blender.org, 1/16/2023) as references for creating a detailed, fully rigged (i.e. articulated) 3D model. (Fig. 3B) The Blender model’s accuracy was checked by using it to create and measure the leg and body dimensions of a 3D printed physical model to confirm that they agreed with those measured for actual SLF nymphs. Next, the 3D model for each specific SLF nymph posture was created by treating the body as a rigid object while configuring the legs to agree with the joint angles observed for the live insect on video. Each 3D model was used to create an .stl file that contains the triangular mesh describing the surface geometry of the model corresponding to each posture.

**Figure 3.**
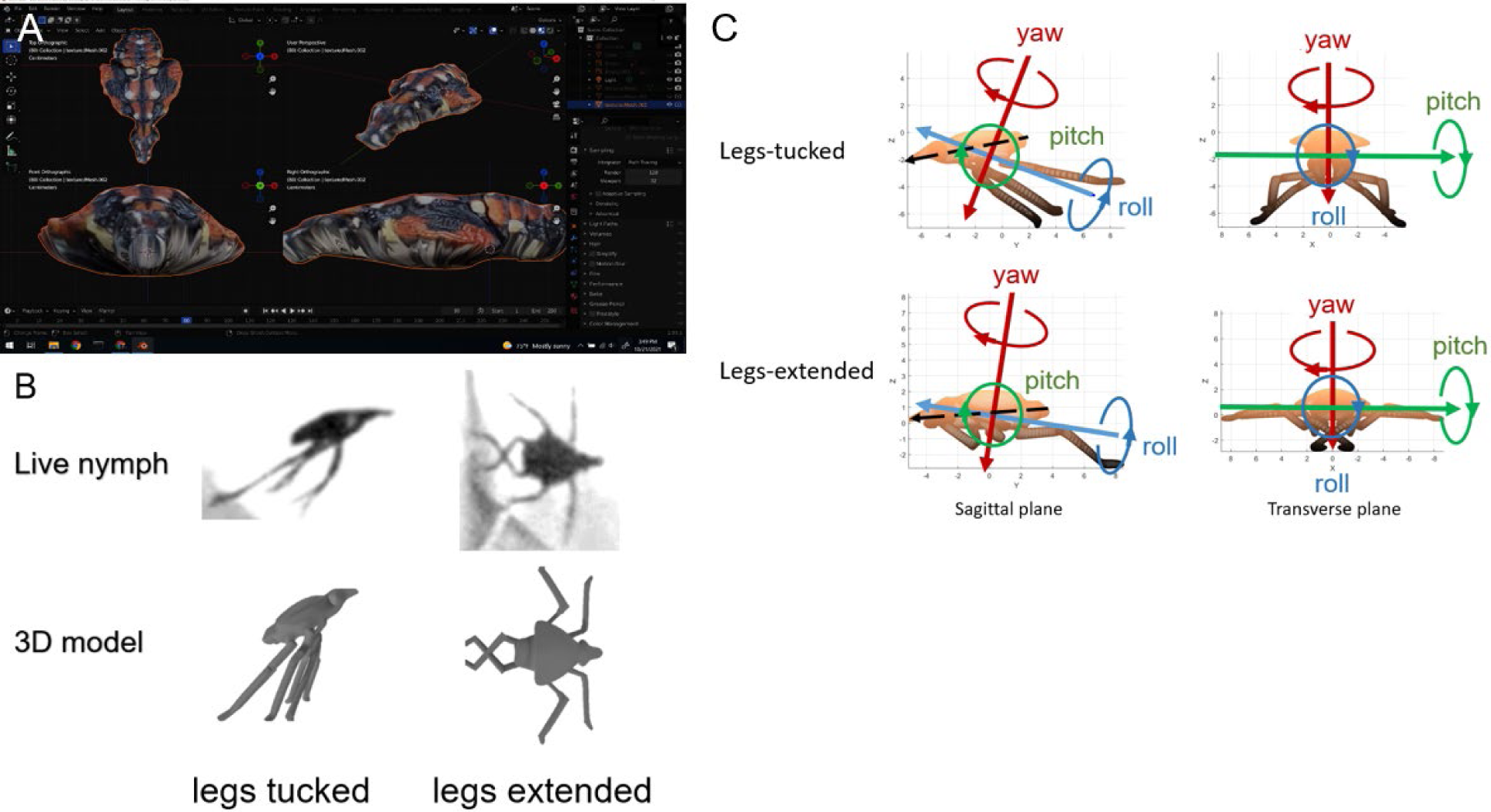
A) Photographs of specimens taken from multiple views were used with photogrammetry software to create detailed 3D renderings of spotted lanternfly nymph bodies and legs. B) Video images of a live spotted lanternfly nymph and the 3D models for two stereotyped midair postures observed during jumping. C) Orientation of the principal axes for the two 3D models, showing the definition of the roll, pitch and yaw axes used in this study.

The MATLAB package RigidBodyParams (Semechko, 2023) was used to compute the volume and COM coordinates for each 3D model, assuming constant density and a body length, *L_b_* = 8.9 mm and mass 28.4 mg typical of 3rd instar SLF specimens. As a check on the accuracy of the 3D model in describing the insect’s body morphology, we used the model’s volume to computed an effective body density 1.0 g/cm^3^, which agrees with published values for insects (Kühsel et al., 2017). The values of **I_*rot*_** = (*I_1_*, *I_2_*, *I_3_*) and the principal axes used for solving Eqn 4 also were computed using RigidBodyParams and 3D models of SLF nymph postures observed on video during different phases of jumping. (Fig. 3C) To interpret how drag forces might vary with posture and orientation relative to airflow, we used ImageJ (Schneider et al., 2012) to find the planar cross-sections of each model (Fig. S2), which correspond to the drag reference areas for intermediate to high Reynolds number (Vogel, 2020).

Because this study was designed to understand the effect of the major changes in posture observed (e.g., extent and angle of leg extension), we used one 3D model for each stereotyped posture observed in our calculations and simulations. SLF 3^rd^ and 4^th^ instar nymphs share a similar body morphology and their masses scale approximately isometrically with body length, *L_b_* (Bien et al., 2023); these 3D models and associated mechanical properties therefore should be reasonable approximations to both life stages when scaled appropriately.

## Results

### Jumping experiments

We recorded analyzable videos of jumping for 83 total trials with successful landings. (Table 1) In addition, we filmed 15 trials of jumping that did not show a successful landing (3^rd^ instars: N = n = 1; 4^th^ instars: N = 13, n = 14). The ethogram in Fig. 4A summarizes the phases and associated stereotyped postures observed during jumping behaviors by SLF nymphs: 1. take-off (Fig. 4B): the insect first assumes a crouching posture, then rapidly extended its hindlegs to provide propulsion, as previously reported for other planthoppers (Burrows, 2009); 2) post-launch (Fig. 4C): the insect keeps its legs tucked close to its body to varying degrees; 3) midair (Fig. 4C): the forelegs and midlegs are fully extended laterally, while the hindlegs are extended rearward and crossed; 4) impact and landing or failure to land: the insect assumes a variety of postures.

**Figure 4.**
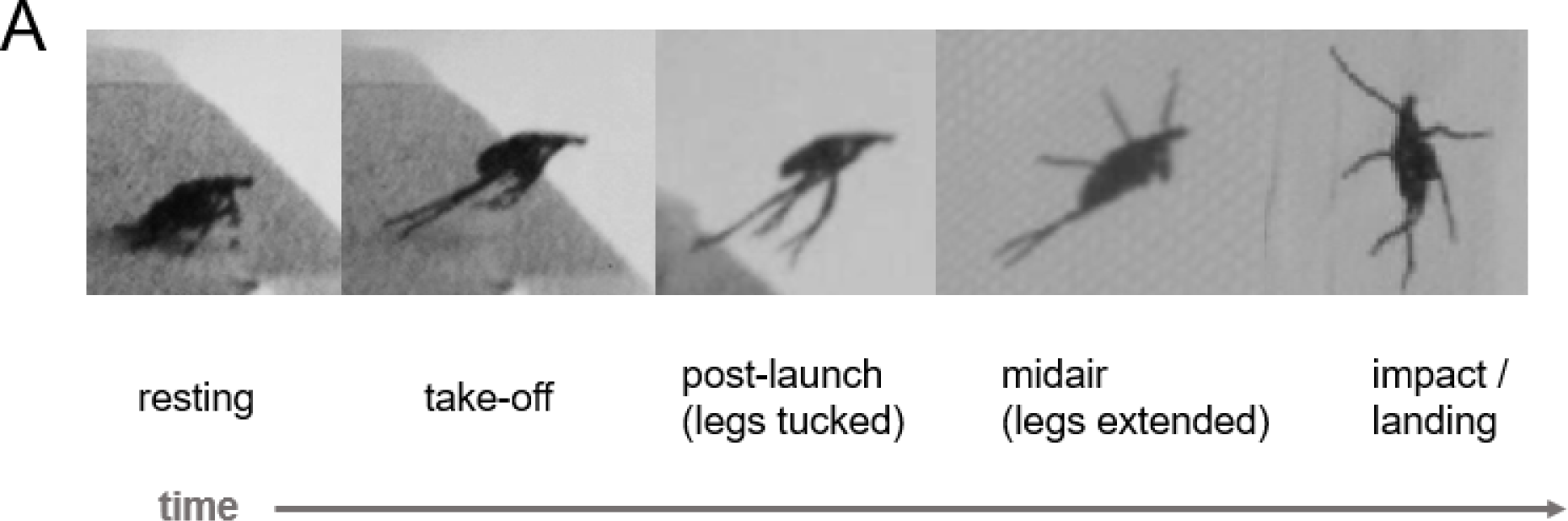
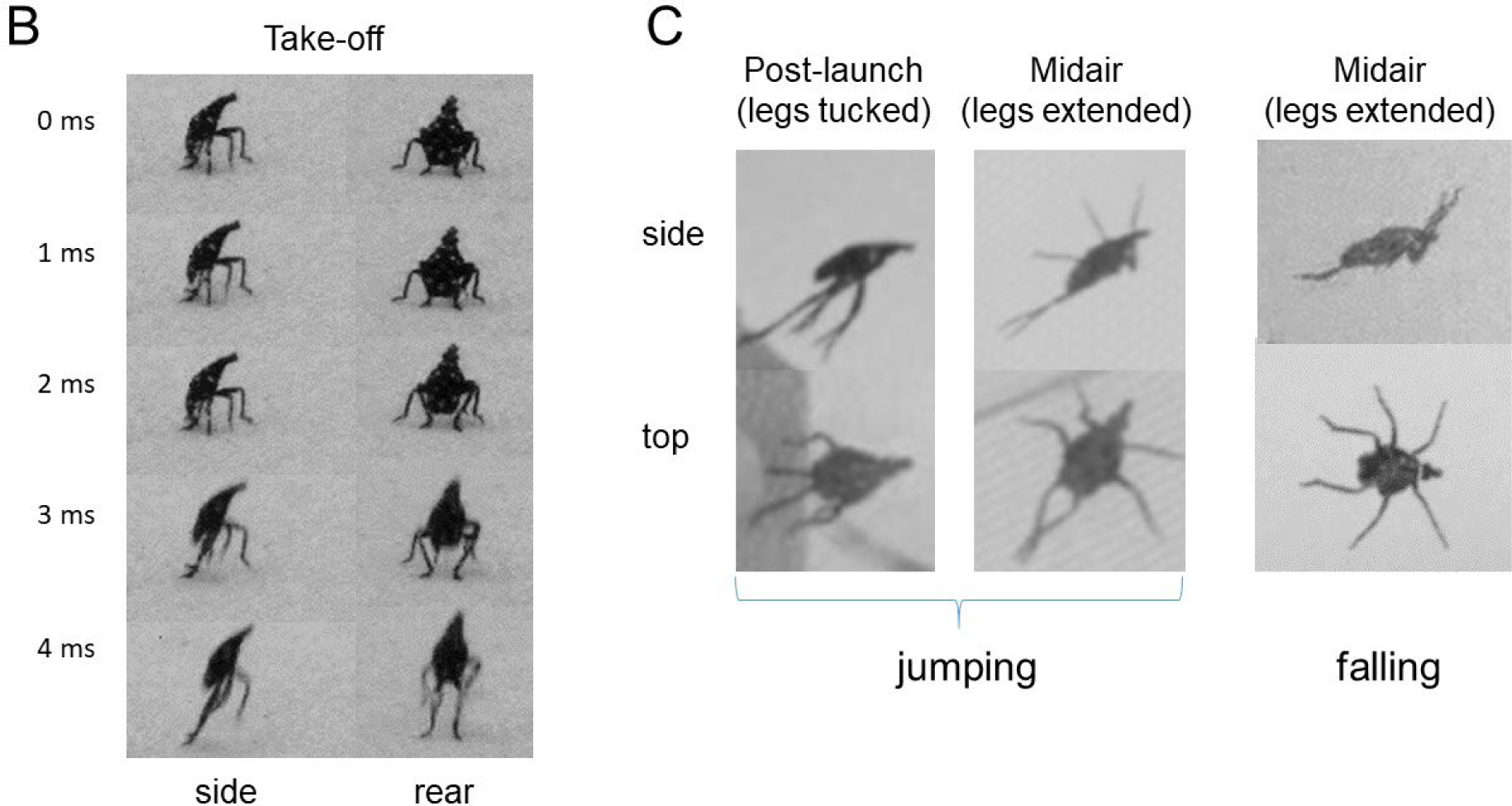
(A) Ethogram of spotted lanternfly nymph jumping behavior divided into observed phases and stereotyped postures. (B) Leg motions during take-off. (C) Stereotyped postures assumed in midair during jumping (this study) and falling (Kane et al., 2021).

**Table 1.** Summary statistics for spotted lanternfly (SLF) nymph jumping kinematics. *M* = mass, *v*_TO_ = take-off speed, *θ* = take-off angle, *T_accel_* = time to reach *v*_TO_, *a*_TO_ = *v*_TO_ / *T_accel_*; N = number individual specimens; n = number of trials, *ω_max_* = maximum rotation rate. All data for SLF nymphs are given as grand mean ± s.e.m (maximum value). * Ranges of values for these parameters reported for a variety of planthopper species studied previously (Burrows, 2009; Burrows, 2014a; Burrows, 2014b; Burrows et al., 2019)).

| Specimen | $M$ (mg) | $v_{TO}$ (ms<sup>-1</sup>) | $\theta$ (deg) | $a_{TO}$ (ms<sup>-2</sup>) | $T_{accel}$ (ms) | $\omega_{max}$ (Hz) |
| --- | --- | --- | --- | --- | --- | --- |
| SLF 3<sup>rd</sup> instar: N = 15, n = 23 | $30 \pm 8$ | $2.7 \pm 0.2$ (5.6) | $35 \pm 4$ (56) | $808 \pm 65$ (1408) | $3.4 \pm 0.1$ | 13 - 53 |
| SLF 4<sup>th</sup> instar: N = 34, n = 60 | $56 \pm 18$ | $3.4 \pm 0.2$ (7.0) | $30 \pm 3$ (58) | $1007 \pm 70$ (2253) | $3.7 \pm 0.1$ | 8 - 45 |
| Other planthopper species* | 2 - 476 | 2.2 – 5.5 | 37 - 69 | $\geq 1060$ | 1 - 5 | 43 |

### Kinematics of the jumping trajectories

Fig. 1C shows plots typical of the measured 3D jumping trajectory coordinates for the entire trajectory and the corresponding zero drag force ballistic fits to Eqn 1 for all 83 trials showing successful landings. The trajectory from take-off to impact was well-described by this model with no contributions from drag or steering: 1) the values of R-squared were consistently high (≥ 0.96); 2) the mean RMSE from the in-plane ballistic fit (0.39-0.43 mm) and <*e_plan_*> for the best fit plane (0.45-0.48 mm) were both close to the 3D reconstruction error (0.54 mm). (See Table S2 for full fit statistics) We therefore used fits to this model performed over the first 25 ms after take-off to characterize SLF nymph jumping performance kinematics. (Table 1) The fitted take-off speeds and angles were used to estimate maximum jumping ranges for comparison with earlier research. For 4^th^ instars, the estimated range of 89 cm agreed with measured values of 86 ±16 cm (this study) and 84 ± 5 cm (Nixon et al., 2021), while the estimated range for 3^rd^ instars, 66 cm, was greater than 45.7 ± 2.8 cm found in (Nixon et al., 2021). The Reynolds numbers (9.3 × 10^2^ ≤ *Re* ≤ 6.6 × 10^3^) computed from the take-off speeds fell within the intermediate regime (10^2^ ≤ *Re* ≤ 10^4^) found for other jumping and flying insects (Bennet-Clark, 1980; Ellington, 1991).

As an additional test for evidence of steering, the difference in the total fit residuals was computed before and after leg extension, and after leg extension and immediately before impact, to look for patterns that would indicate changes either in direction or in angular velocity. No significant differences were found in either case. (Table S3)

Shortly after take-off, all but one of the SLF nymphs filmed assumed a stereotypical midair posture (“legs-extended”) in which the forelegs and midlegs were extended in the transverse plane and the hindlegs crossed and extended rearward. (Table 1; Movie 1) The time from take-off to full leg extension, *T_ext_*, was 40 [36, 44] ms for 3^rd^ and 48 [45, 51] ms for 4^th^ instar nymphs. Specimens maintained this crossed-leg posture even after impact and landing in 12/23 trials for 3^rd^ and 21/61 trials for 4^th^ instar nymphs. (Movie 1) No other typical changes in posture were observed during the midair phase of jumping.

Rotational kinematics

All analyzable videos recorded in the laboratory (n =98) and field (n = 6) showed specimens rotating after takeoff with a variety of initial angular velocity directions and subsequent rotational motions. The angular speed of the cranial-caudal axis, *ω_cc_*, vs time was tracked and analyzed for a total of 21 arbitrarily-selected trials. (Fig. S3) After an initial analysis found no statistical difference between the rotational kinematic measures for 3^rd^ and 4^th^ instars, the results were pooled across life stages. (Table S4) To provide an estimate of the rotational drag timescale to use in simulations, the data for *ω_cc_*, vs time were also fit to Eqn 5b, giving *T*_*drag*_ = 155 [137, 174] ms for the legs-tucked and 76 [60, 92] ms for the legs-extended postures. (Fig. S3, Table S4) (We fit to a single value of *T*_*drag*_ for each posture because the orientation of the rotating nymphs constantly changed relative to the airflow direction.)

Next, the estimated moments of inertia, *I_i_*, and cross-sectional areas (Table 2) and principal axis directions (Fig. 3C) were computed for the legs-tucked and legs-extended 3D models for use in rotational dynamics simulations. While the transverse (pitch) axis agrees with the **û_2_** principal axis for both 3D models, the other two principal axes do not correspond with the body cranial-caudal and dorsal-ventral axes. We therefore refer to rotations about principal axes **û_1_** with moment of inertia *I_1_* as roll, and **û_3_** with moment of inertia *I_3_* as yaw, because they most closely agreed with the expected anatomical axes for these rotations. The disagreement between the anatomical axes and principal axes for roll and yaw means that the measured angular value of *ω_cc_* depends on roll, pitch and yaw rotations.

**Table 2.** The moment of inertia for each whole-insect 3D model created for the midair postures observed during jumping by spotted lanternfly nymphs.

| | Moment of inertia, $I \pm 0.1 (\times 10^{-10} \text{ kg m}^2)$ | | | Drag reference area $\pm 1.5 (\text{mm}^2)$ | | |
| --- | --- | --- | --- | --- | --- | --- |
| Posture | Roll | Pitch | Yaw | Transverse | Dorsal | Sagittal |
| Legs-tucked | 2.28 | 4.60 | 5.30 | 22.6 | 51.7 | 35.7 |
| Legs-extended | 2.89 | 3.28 | 5.48 | 29.2 | 54.6 | 21.4 |
| Tucked/<br>Extended | 0.79 [0.75, 0.83] | 1.40 [1.35, 1.45] | 0.97 [0.94, 1.00] | 0.77 [0.67,0.87] | 0.95 [0.87, 1.03] | 1.48 [1.41, 1.55] |

Taken together with the unequal values of *I_i_* computed for both models, this means that the direction of the axis of rotation should vary in time unless the insect manages to take-off rotating purely around a single principal axis. In fact, most plots of the measured *ω_cc_* vs time displayed oscillations due to angular momentum twisting in which the nymph’s motion varied between different combinations of roll, pitch, and yaw rotations. (Fig. 5, S4; Movie 1) The mean period for these oscillations was *T_osc_* = 36 [31, 41] ms for all analyzed tracks, independent of the value of *ω_TO_* (linear regression: R-squared = 0.05, F-statistic = 0.85, P = 0.37).

**Figure 5.**
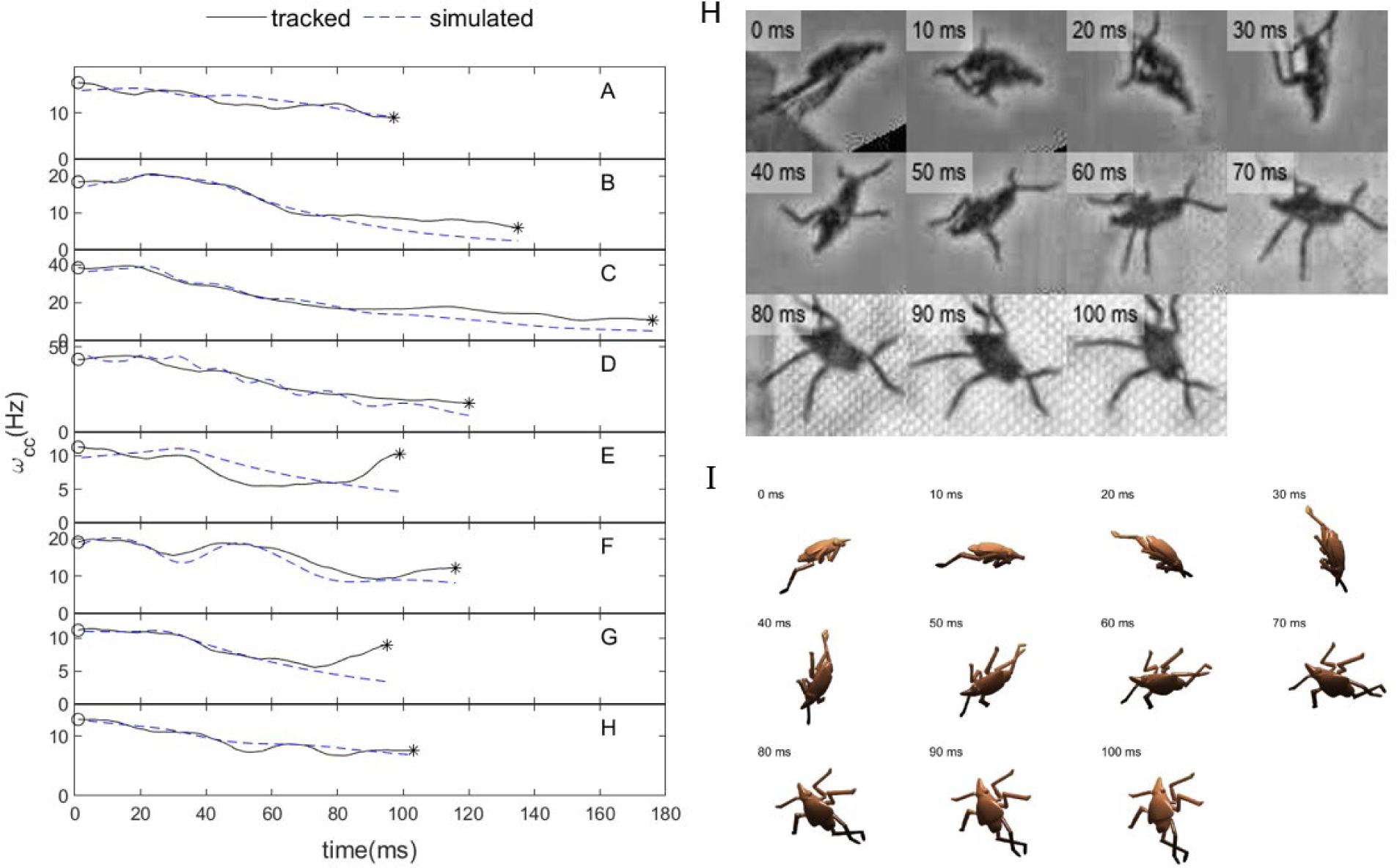
(A)-(G) Angular velocity of body orientation (cranial-caudal axis), *ω_cc_*, vs time tracked on video and from simulations using parameters measured from the tracked data. Details of the simulations are described in the Methods. (Symbols: ○ = take-off; * = impact) H) Image sequence from the video H) and matching simulations I) for the data in E; the SLF model is shown in the legs-extended posture for all frames.

To interpret these results, we used Eqn 4 and 5a to simulate the rotational dynamics of SLF nymphs during the midair phase of jumping for n = 8 representative trials for which rotational kinematics were tracked and for which we were able to determine the angular velocity vector at take-off. For the interval of time, *T_ext_*, over which the moment of inertia changed due to leg extension, ***I_rot_***(*t*) was estimated by linearly interpolating the inertia tensor along each principal axis between those corresponding to each posture’s 3D models using:

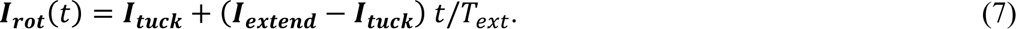

The resulting time-dependent inertia tensor was used to estimate effects due to posture change due to changes in the principal axis directions and values of *I_1_*, *I_2_*, *I_3_*. The simulated values of *ω_cc_* vs time were then averaged over 25 ms for comparison with the video analysis. (Fig. 5) Results of the Wilcoxon two-sample signed-rank test for paired values of tracked and simulated *ω_cc_*(t) indicated that the measured and simulated data did not differ significantly (p < 0.014). (Table S6) The measured and simulated data exhibited a similar overall functional behavior (i.e., a decrease in *ω_cc_* with increasing time modulated by a lower amplitude oscillatory component). The oscillation period measured from simulations (*T_osc_* = 32.0 [23.3, 40.8]) agreed with tracked values (29.4 [21.7, 37.1]ms for the simulated datasets (Wilcoxon ranked sum, W = 7.0, p = 0.63, n = 5) although the simulations did not reproduce the amplitude and phase of the oscillations accurately in most trials analyzed. They also were unable to account for the abrupt increase in angular velocity before impact observed in 6 of 21 trials. (Fig. S3)

Landing behavior

In spite of their rotational behavior in the air, SLF nymphs were filmed landing securely on vegetation (outdoors) or on the target surface (in the laboratory). (Movie 1) A total of 207 videos filmed in the laboratory showed the body orientation relative to the surface on impact. Landing outcome data for the 3^rd^ and 4^th^ instar nymphs were pooled because the distributions of impact orientation were not significantly different (chi-squared two-sample test: χ^2^ = 24, p = 0.24 successful landings; χ^2^ = 18; p = 0.26 failed landings). Significantly more specimens landed successfully than failed (probability of successful landing 58.9% [51.9, 65.7]%; binomial confidence interval, Clopper-Pearson method, N = 207). While specimens were observed landing successfully after impacting the surface at all possible orientations, successful landings were observed significantly more often than expected for impacts on the ventral side, and failed landing attempts were significantly more frequent than expected for dorsal impacts. (Fig. 6A,B) All other landing outcomes had no significant dependence on orientation. For all outcomes after impact, the ventral orientation was the most frequent and occurred significantly greater than expected. (Fig. 6C)

**Figure 6.**
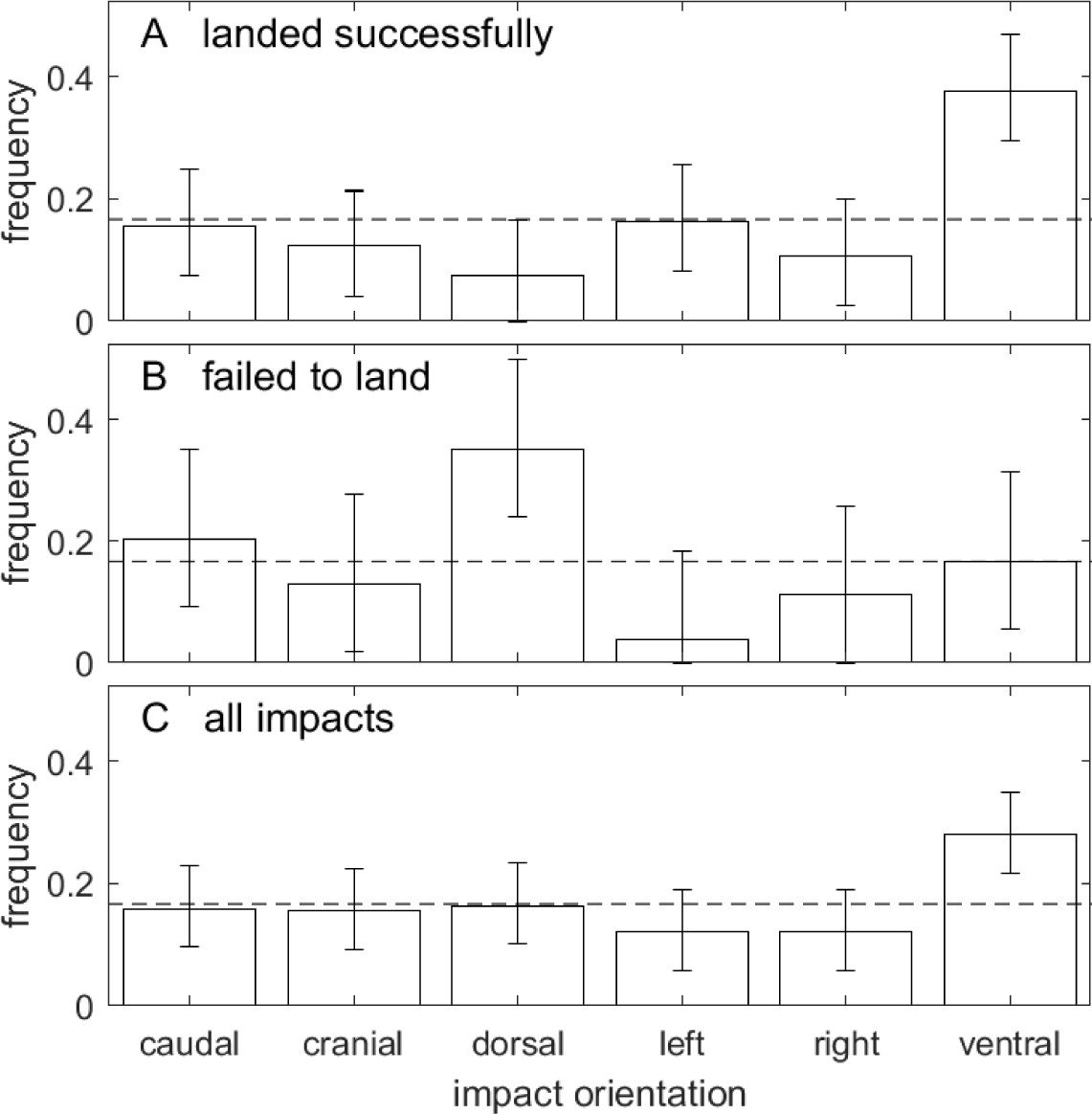
Spotted lanternfly nymph body orientations at impact for all A) successful landings (n = 122) and B) failed landing attempts (n = 85). The error bars are 95% CI for each measured proportion. The dashed line at 1/6 corresponds to equal probability of landing in each orientation.

While sometimes the SLF nymphs were able to land securely upon first impact, there were multiple instances of them initially grasping the surface by only one or two tarsal claws and struggling to achieve a secure hold. (Movie 1, Fig. S4) The compliant fabric target appeared to absorb kinetic energy and allow the insects to grasp hold using their tarsal claws. A total of 25% (31/122) videos showed specimens bouncing off the surface entirely at first impact but landing on a second try, either by sliding down the initial surface while trying to grasp it repeatedly or by bouncing onto and grasping another surface before reaching the ground. Some specimens performed a bellyflop landing; i.e., they first impacted on a side or the cranial or caudal end, then pivoted onto their ventral surfaces.

## Discussion

The invasive spotted lanternfly is difficult to eradicate in part due to its agile jumping abilities. Our study revealed that SLF nymphs regularly perform acrobatic jumps with complicated, but controlled, aerial rotations in the field and laboratory environments, with a high rate of landing successfully. In spite of their fast rotational motion, their jump trajectories agreed with zero-drag ballistic motion with performance measures similar to those reported for other planthoppers, providing no evidence of active control of their trajectory direction. Using photogrammetry, 3D modeling, and simulation, we were able to reconstruct the complex, 3D rotational movements during a jump. This analysis also demonstrated how limb extension and subtle changes in limb position dramatically alter the moment of inertia and the principal axes of rotation, permitting fine adjustments to their body orientation.

While in the air, SLF nymphs were found to spin rapidly (5-45 Hz) about rotational axes that varied in time, indicating their motions indeed correspond to angular momentum twisting rather than pure roll, pitch or yaw rotations about a single body axis. The main posture adjustments observed during jumping involved holding the legs tucked to varying degrees after take-off and then extending them rapidly 40-48 ms after take-off, comparable to measured insect neuromuscular response times (Sponberg and Full, 2008). Calculations using 3D modeling showed that this leg extension corresponds to “throws” of body appendages (i.e., changes in both the moment of inertia and in the body’s principal axes of rotation) that led to complex changes in subsequent rotational motion.

Using only parameters based on the angular velocity at take-off, two simplified 3D models of jumping postures, and drag torques based on fits to tracked orientation data, we were able to simulate the measured body orientation’s angular speed, *ω_cc_*, vs time data so as to reproduce its overall dependence on time, the 11% decrease in *ω_cc_* after the legs were fully extended, and oscillations in *ω_cc_* due to the time-varying direction of the axis of rotation. The simulations showed that the rotational dynamics model needed to include the effect of aerodynamic drag torques to correctly describe the observed near-monotonic decrease in angular velocity during the midair phase. Modeling the legs’ configurations accurately using the 3D models was found to be important because they accounted for 61-82% of the moment of inertia values, enabling leg extension to cause large changes in the moment of inertia and principal axis of rotation, and therefore in both the rotational speed and axis direction. (Table 2, S5) In combination with Eqn 7b, this indicates that, for example, small pitch rotations of the legs result in a 3.83× greater change in body orientation in the legs-extended posture, providing the nymph with the ability to make attitude adjustments with fine corrections to the extension and orientation of its legs. Therefore, SLF nymphs can adjust their axis of rotation, and hence body orientation, by flexing, extending or rotating their legs to achieve attitude control.

In spite of their rotational motion, SLF nymphs were observed to be able to land securely in 58.9% of impacts. The finding that specimens impacted on their ventral sides significantly more often than predicted supports their use of attitude control. The results from rotational dynamic simulations indicates that their observed body posture changes likely contribute to this ability. In addition, non-streamlined objects moving in the SLF nymph’s aerodynamic regime tend to orient with the maximum area normal to the airflow direction (Bennet-Clark, 1980; Vogel, 2020). Because our 3D models found the dorsal plane has the maximum area during both jumping postures, this suggests that aerodynamic drag torques tend to orient the body to the most likely orientation for successful landing, as previously reported for various insect species during falling (Ribak et al., 2013; Yanoviak et al., 2010; Zeng et al., 2017) and jumping (Burrows et al., 2015). (Table 2)

While landing attempts were disproportionately successful for impact on the ventral surface, SLF nymphs were able to land securely for all orientations at impact. They did so by grasping the surface by one or more tarsi while rotating rapidly and bouncing, similar to the capabilities observed for falling SLF nymphs landing on leaves (Kane et al., 2021) and for frogs landing on sticks (Bijma et al., 2016). This ability is of particular importance in their natural environment, where foliage landing spots present themselves at all orientations and locations, the ground is likely covered with clutter, and many landing targets recoil on impact. Thus, landing success can be enhanced by a number of behaviors, including achieving a preferred orientation when possible, but also by negotiating the aftermath of collisions so as to cling to the target (Jayaram et al., 2018; Kane et al., 2021; Reichel et al., 2019; Siddall et al., 2021).

The active control of body posture during jumping also serves functions other than rotation and attitude control. For example, SLF nymphs displaced by predators or disturbances benefit from landing on or near their current host plant in order to retain protective cover, avoid expending energy on finding and climbing a new host, and resume feeding. This is facilitated by extending their legs to increase the likelihood of encountering leaves while in the air. To estimate this effect, consider that the cross-section for encountering an obstacle can be estimated from the area of the minimum polygon enclosing the relevant body plane; these mean areas are 130 and 180 mm^2^ for the legs-tucked and -extended models, respectively, giving a 38% increase in area due to leg extension. Furthermore, because these insects often land using the same bellyflop method observed for frogs landing on sticks (Bijma et al., 2016), extending their legs to facilitate this landing method also could dissipate energy on impact to help stabilize landings. Furthermore, the midair posture during jumping differs from that observed for falling arthropods (Kane et al., 2021; Ribak et al., 2013; Yanoviak et al., 2015) in that SLFs, like planthoppers, leafhoppers, treehoppers, fleas and locusts (Burrows, 2009; Burrows, 2011; Burrows, 2013; Burrows, 2014a; Burrows, 2014a; Burrows and Dorosenko, 2017; Burrows and Sutton, 2008; Burrows et al., 2007), cross their hindlegs in the leg-extended pose. Since the crossed hindlegs function as a single unit on impact, this configuration also might provide protection against buckling on impact (Burrows and Sutton, 2008). Given that SLF nymphs were able to keep their hindlegs crossed even after impact, it would be interesting to investigate their leg morphology for a latching mechanism similar to the interlocking hairs or gear-like mechanism that some planthoppers use to synchronize leg motion (Burrows and Sutton, 2013).

These finding suggest multiple directions for future work. During take-off, a net torque can result from asymmetrical force production by the hindlegs (Burrows and Sutton, 2013) or from extending the body COM forward of the hindlegs push-off surface (Burrows, 2012). This indicates that rotations are likely for insects taking-off from flexible stems and leaves with varying orientations, as observed in our field videos. Consequently, it would be interesting to investigate how taking-off and landing behaviors differ for leaves, branches and solid ground with differing compliance and textures (Burrows and Sutton, 2008; Mo et al., 2020a; Ribak et al., 2012), as well as the effect of wind speed and orientation (Aguilar-Arguello et al., 2021; Wolfin et al., 2019).

Moreover, while field studies of SLFs have found that adults tend to land on vertical, high contrast structures such as posts (Baker et al., 2021; Wolfin et al., 2019; Zeng et al., 2020), this possibility has not been investigated for nymphs. While SLF nymphs did not show evidence of steering in this study, they were orienting toward a uniform target a relatively short distance (0.21 ± 0.06 m (mean ± sd)) away compared to their mean ranges. Given that many other insects are known to steer toward preferred landing targets (e.g., leafhoppers (Brackenbury, 1996), flea beetles (Brackenbury and Wang, 1995) and stick insect nymphs (Zeng et al., 2015)), future work should consider the trajectories of jumping in naturalistic settings to see if these insects exhibit targeted midair descent over their entire jumping ranges.

While the results of this study demonstrate the utility of 3D modeling for use in simulating rotational dynamics in biomechanics, we only considered two characteristic postures observed during jumping by SLF nymphs and transitions between these postures. The models presented here therefore were not intended to capture the effect of differences between individual body morphology or smaller changes in posture due to finer-scale leg adjustments during jumping that may also influence attitude control. In future work, it would be interesting to model the effect of leg motions in greater detail. One approach would be to obtain higher resolution video images of the insects to allow tracking of multiple body keypoints to determine body posture and 3D orientation in detail. Another is to perform simulations using realistic 3D rendered models to inform the construction of articulated models with accurate mechanical properties but more easily-simulated geometries (e.g., with cuboidal bodies and jointed rods for legs and appendages), similar to the jointed-rod model used successfully in earlier work (Burrows et al., 2015; Zeng et al., 2017).

The interplay between the complex jumping behavior of insects can yield new insights for animal behavior and for bioinspired jumping robotics. The approaches used here provide new insights into the physical mechanisms at work during aerial reorientation, and can easily be generalized to other organisms and motions.

## Acknowledgements

We would like to thank Robert Beyer for help in designing and fabricating the experimental apparatus.

## Data availability

The code and datasets supporting this article have been uploaded as part of the supplementary material as Dataset S1.

### List of symbols and abbreviations

SLF: spotted lanternfly
3D: three-dimensional
COM: center of mass
*v_TO_*: take-off speed
*a*_TO_: acceleration during take-off
*T_accel_*: acceleration time for take-off
*θ*: take-off angle relative to horizontal
*e_ballistic_*: ballistic fit residuals
*e_planar_*: planar fit residuals
*R*: maximum jumping range
*T_TOF_*: time-of-flight
*Re*: Reynolds number
*ρ*: air density
*L_b_*: body length
*μ*: dynamic viscosity of air
*τ*: torque (moment of force)
*r*: moment arm magnitude
*F*: force
*L*: angular momentum
*I_rot_*: moment of inertia
*Ii*: moment of inertia along the ith principal axis
*û_i_*: unit vector lying along the ith principal axis
*ω*: angular velocity
*T_drag_*: characteristic rotational drag timescale
*ω_cc_*: angular speed of the cranial-caudal axis
*ω_TO_*: *ω_cc_* immediately after take-off
*T_flip_*: characteristic time for flipping (intermediate axis theorem)
*T_osc_*: oscillation period

